# The actin cytoskeleton of the mouse sperm flagellum is organized in a helical structure

**DOI:** 10.1101/252064

**Authors:** María Gracia Gervasi, Xinran Xu, Blanca Carbajal-Gonzalez, Mariano G Buffone, Pablo E Visconti, Diego Krapf

## Abstract

Conception of a new mammalian organism is determined by the fusion of a sperm cell with an oocyte during fertilization. Motility is one of the features of sperm that allows them to succeed in fertilization, and their flagellum is essential for this function. Longitudinally, the flagellum divides into the midpiece, the principal piece, and the end piece. A precise cytoskeletal architecture of the sperm tail is key for the acquisition of fertilization competence. It has been proposed that the actin cytoskeleton plays essential roles in the regulation of sperm motility, however, actin organization in sperm remains elusive. In the present work, we found different types of actin structures in the sperm tail, using stochastic optical reconstruction microscopy (STORM). In the principal piece, actin is radially distributed between the axoneme and the plasma membrane. The actin-associated proteins spectrin and adducin are also found in these structures. Strikingly, polymerized actin in the midpiece forms a double-helix that accompanies mitochondria. Our findings illustrate a novel specialized structure of actin filaments in a mammalian cell.

---

Mammalian sperm are highly polarized and specialized cells composed of a head and a tail with a single, well-defined goal: to fertilize an oocyte (Yanagimachi, 1994). This specific functionality is granted by a tightly organized cellular structure. The sperm tail and head are joined by a connecting piece and are further organized into different compartments with specific functions in motility, ATP generation, exocytosis and sperm-egg fusion (Eddy, 2006). Some of these subdivisions are membrane-enclosed and others are separated by diffusion barriers that segregate proteins and metabolites. Longitudinally, the sperm tail is divided into the midpiece, the principal piece and the end piece. The midpiece and the principal piece are separated by the annulus, a septin-rich ring structure. Radially, these compartments are arranged around the axoneme, which consists of a central pair surrounded by nine peripheral microtubule doublets (Lindemann and Lesich, 2016). In both the midpiece and the principal piece, the axoneme is surrounded by outer dense fibers (ODFs). In the midpiece, the ODFs are wrapped in mitochondria organized in an end-to-end touching configuration that forms a double helical structure (Otani et al., 1988). On the other hand, the principal piece ODFs are surrounded by a fibrous sheath (Lindemann and Lesich, 2016). The midpiece and principal piece structural organizations are essential for sperm function. Sperm presenting defective structures in either the mitochondrial sheath, the fibrous sheath, or the outer dense fibers have abnormal morphology as well as impaired motility and fertility (Bouchard et al., 2000; Chemes et al., 1987; Kissel et al., 2005; Miki et al., 2002; Olson et al., 2005).

The scaffolding framework that maintains the organization of the sperm flagellum cytoskeleton and its differentiated structures is poorly understood. In addition to the axoneme, actin and actin-associated proteins have been documented in sperm from many species (Romarowski et al., 2016). Actin is an abundant and highly conserved protein among eukaryotes, that is essential for diverse cellular functions such as cell shape, motility, membrane organization, and cytokinesis (Campellone and Welch, 2010). Diverse proteins interact with actin filaments and form complex cytoskeletal structures that ensure cell functionality. In erythrocytes, polymerized actin together with spectrin and adducin form a sub-cortical meshwork that gives the cell membrane the elasticity required to survive in the circulatory system (Lux, 2016). In neurons, short F-actin filaments capped by adducin are organized as periodic rings interconnected by spectrin along the axon (Xu et al., 2013). In HEK cells, a cortical actin meshwork forms a fractal structure that organizes the plasma membrane (Andrews et al., 2008; Sadegh et al., 2017). Actin has been reported in different compartments of mammalian sperm including the tail (Breed and Leigh, 1991; Fouquet and Kann, 1992), and actin dynamics have been proposed to play a role in sperm motility and the acrosome reaction (Breitbart et al., 2005; Brener et al., 2003; Itach et al., 2012). However, despite the importance of actin organization and dynamics, little is known about the structure, function and regulation of actin in sperm.

In this article, we use stochastic optical reconstruction microscopy (STORM) to uncover the actin-based cytoskeleton in the sperm flagellum. We find that the sperm midpiece and principal piece depict specialized actin structures. In the midpiece, polymerized actin forms a double-helix that follows the mitochondrial sheath, a type of filamentous actin structure that has not been observed so far. On the other hand, actin in the principal piece is radially distributed from the axoneme to the plasma membrane. In addition, the actin-associated proteins spectrin and adducin, which are involved in shaping the actin cytoskeleton in other cell types, are also organized differentially in both sperm tail compartments. These results suggest a role of polymerized actin in the molecular organization of flagellar structures involved in motility regulation and sperm function.

## Results

### F-actin localization and structure in mouse sperm midpiece

The localization of F-actin in mouse sperm was initially determined by immunofluorescence using two different approaches: i) labeling with fluorescent phalloidin, and ii) using a transgenic mouse line that expresses LifeAct-GFP, a small peptide that stains F-actin without altering its dynamics, in all cell types (Riedl et al., 2010). In both cases, F-actin was observed in the sperm head and throughout the tail, with higher intensity in the midpiece (Fig. 1 A and B). This localization is in line with previous reports of F-actin in sperm (Bouchard et al., 2000; Flaherty et al., 1986; Romarowski et al., 2015); however, the narrow shape of the sperm flagellum together with the limitations in resolution of widefield microscopy, did not allow to observe details of the actin structure. To investigate the organization of F-actin in the flagellum in detail, we imaged phalloidin-stained sperm using STORM (Bates et al., 2008), a superresolution technique that was previously employed successfully to evaluate the localization of different proteins in mouse sperm (Alvau et al., 2016; Chung et al., 2014). Consistent with results obtained using widefield immunofluorescence imaging (Fig 1 A and B), STORM showed high density of F-actin in the tail. In addition, STORM revealed two different structures in the midpiece and the principal piece (Fig. 1 C). While in the midpiece F-actin forms a periodical structure, in the principal piece is uniformly distributed along the flagellar length. Figure 1 D shows a zoom of the periodical arrangement of F-actin found in the sperm midpiece, this structure is also shown in the Supplemental video 1. The cross-section of the midpiece showed that F-actin is completely absent from the central axoneme (Fig. 1 E). It is worth notice that, besides the prevalent helical structure observed in the midpiece, short F-actin bundles are found spread along the circumference of the flagellum (Fig. 1 F and G).

**Figure 1:**
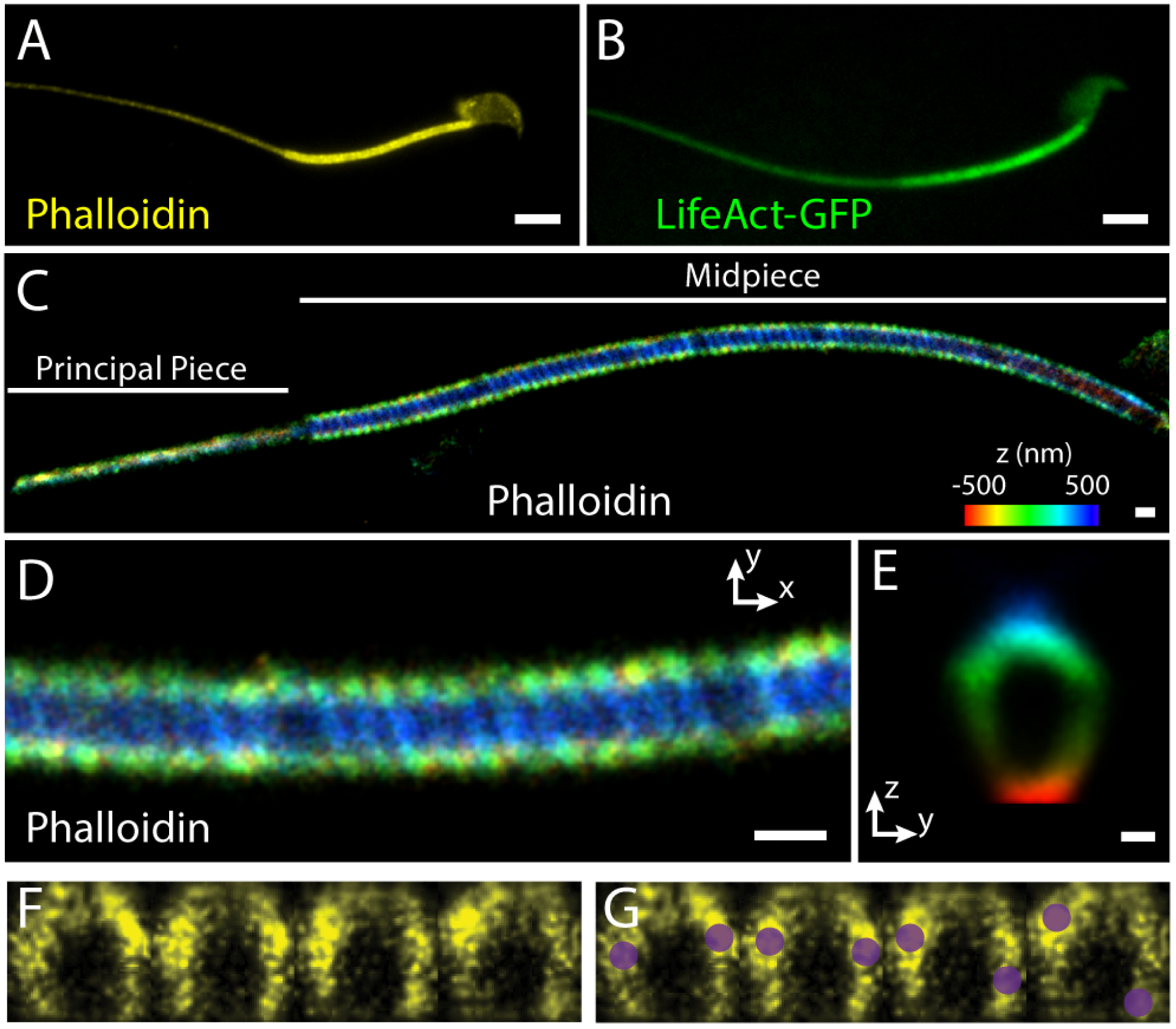
Actin structure in the sperm flagellum. **(A)** Widefield image of actin labeled with Phalloidin-AF647. **(B)** Sperm from transgenic mouse that expresses GFP-LifeAct. (C) STORM reconstruction of actin labeled with phalloidin-AF647 in the sperm tail. The whole midpiece and part of the principal piece are observed in the image as lageled. The reconstruction is colored according to the localization height as indicated in the colormap. **(D)** Zoom of part of the midpiece of the sperm shown in panel C. **(E)** Composite sum of intensity projection of cross-sections along 7 μm of the sperm midpiece. **(F)** Cross-sections of the actin superresolution image separated by 39 nm along the sperm axis. **(G)** Sketch of the location of the higher intensity structure overlaid on the cross sections in panel E. The scale bars in panels A-B are 5 μm, in panel C and D are 500 nm, and in panel E is 200 nm.

In order to exploit the cylindrical symmetry of the flagellum, we converted the molecule localizations as found by STORM into cylindrical coordinates (*r*, θ, *z’*). Note that when working in cylindrical coordinates we refer to *z’* as the flagellum axis but in Cartesian coordinates it is the direction normal to the coverslip, so that in Cartesian coordinates the sperm lies on the *xy* plane, as sketched in Fig. 2 A. Figure 2 B shows the distribution of F-actin radial localizations *P*(**r**) in the midpiece for different individual cells, together with the mean distribution. The distribution of radial localizations is presented as an area density with normalization 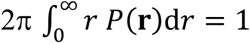 and, thus, the mean radius is obtained as

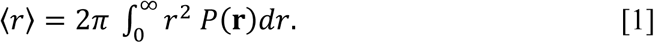

**Figure 2:**
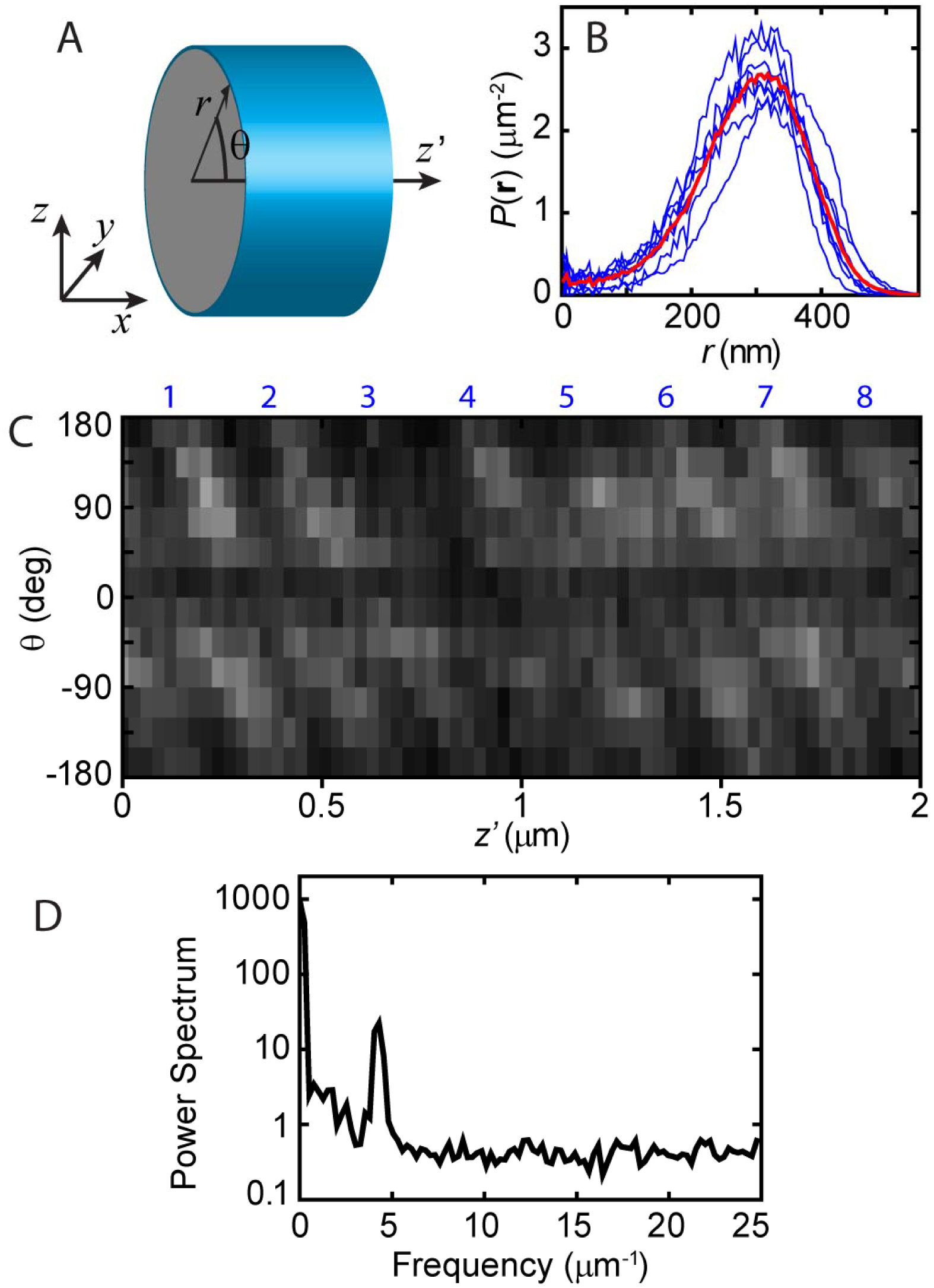
Analysis of the actin structure in the sperm midpiece. **(A)** Sketch of the Cartesian and cylindrical sets of coordinates used in the article in relation to the sperm flagellum geometry. **(B)** Radial distribution of F-actin localization in the midpiece. The blue lines are distributions from seven individual cells and the thick red line is their average. **(C)** Two-dimensional histogram of F-actin localization in terms of azimuth angle *q* and axial position z’ in the sperm midpiece reveals stripes that follow a double helix. The blue numbers on the top of the histogram indicate gyre number as guides to the eye. **(D)** Representative single-cell power spectrum of the actin axial localization. The power spectrum is computed independently for each angle *q* and then all the power spectra from the different angles in a single cell are averaged as described in the text. A peak at 3.92 ± 0.22 μm^-1^ indicates the periodicity of the actin structure in this representative cell.

The radial distribution in the midpiece (Fig. 2 B) indicates that F-actin peaks at ∼ 300 nm from the center of the flagellum. A graphical representation of F-actin localization in terms of the azimuth angle θ vs. axial distance *z’* along the midpiece shows that the periodical F-actin organization found in this region forms a helical structure (Fig. 2 C). It is possible to trace an actin line in this representation from the angle 180° to -180°, at which point the trace connects to a different line at 180° as expected from a helix. However, when connecting these traces one trace is skipped; namely we observe one continuous helix composed of traces 1, 3, 5, … that alternate with a second helix composed of traces 2, 4, 6, …, as labeled on the top of Fig. 2 C. Thus, the actin structure forms two congruent helices. The distance between the F-actin helices was accurately measured cell-by-cell using a Fourier transform method. First, the θ vs. *z’* representation was coarse grained into bins along θ as shown in Fig. 2 C. Each angle θ yields a series of intensities *I*_θ_ (*z*). Then, a power spectrum was obtained from each intensity series by means of a Fourier transform. Using an azimuthal angle bin size of 30°, we obtained 12 different power spectra in each image that were averaged to obtain one spectrum for each cell as shown in Fig. 2 D. In this representative cell, the spectrum shows a single (non-zero frequency) peak at a spatial frequency *f*_0_ = 3.92±0.22 μm^-1^ (peak ± HWHM). This analysis was performed in multiple cells, with all cells yielding a similar peak in the power spectrum. Namely the mean peak corresponds to a distance between helices 1/〈*f*_0_〉 = 243.7 ± 1.8 nm (mean ± sem, n = 15).

Figure 3A shows a transmission electron microscopy (TEM) image of a longitudinal section of the sperm midpiece. The mitochondrial organization, which displays a similar periodicity to the observed actin structure is clearly visible. Thus, the helical actin structure in the midpiece follows the organization of the mitochondrial sheath. The red arrow in the figure indicates a distance of 300-nm from the center of the flagellum, which corresponds to the F-actin radius in the midpiece as found by STORM. Based on these measurements, the helical structure of actin presented a radial localization coincident with the radial center of the mitochondria, suggesting that the actin filaments are located in thin regions between mitochondria. The surface of the midpiece can also be investigated in extremely high resolution by atomic force microscopy (AFM). Figure 3 B shows an AFM topographic image of a sperm midpiece. Helical grooves, with the same periodicity as seen for actin, are also observed to decorate the surface. These measurements established that the midpiece surface exhibits the same pattern of the actin cytoskeleton. From the actin STORM reconstructions, it is possible to count the number of full helical turns (gyres) in the midpiece. We found that each of the two actin helices makes 43.7 ± 0.25 (n = 6 cells) full helical turns along the midpiece which makes a total of 87 turns when considering both helices. This number is in excellent agreement with the number of mitochondria windings around the flagellum. On average, the mouse sperm midpiece comprises a total of 87 mitochondria gyres arranged in a double helix (Eddy, 2006). We further investigated the organization of F-actin and the mitochondrial sheath by two-color structural illumination microscopy (SIM), a technique that allows an axial resolution of ∼ 120 nm. Figures 3 C-E show the dual imaging of mitochondria labeled with MitoTracker Green FM and actin stained with Alexa Fluor 647-conjugated phalloidin. Consistent with our STORM results, the periodical structure of F-actin was also observed by SIM. Most importantly, the localization of F-actin appeared to generally alternate with the mitochondria arrangement (Fig. 3 E).

**Figure 3:**
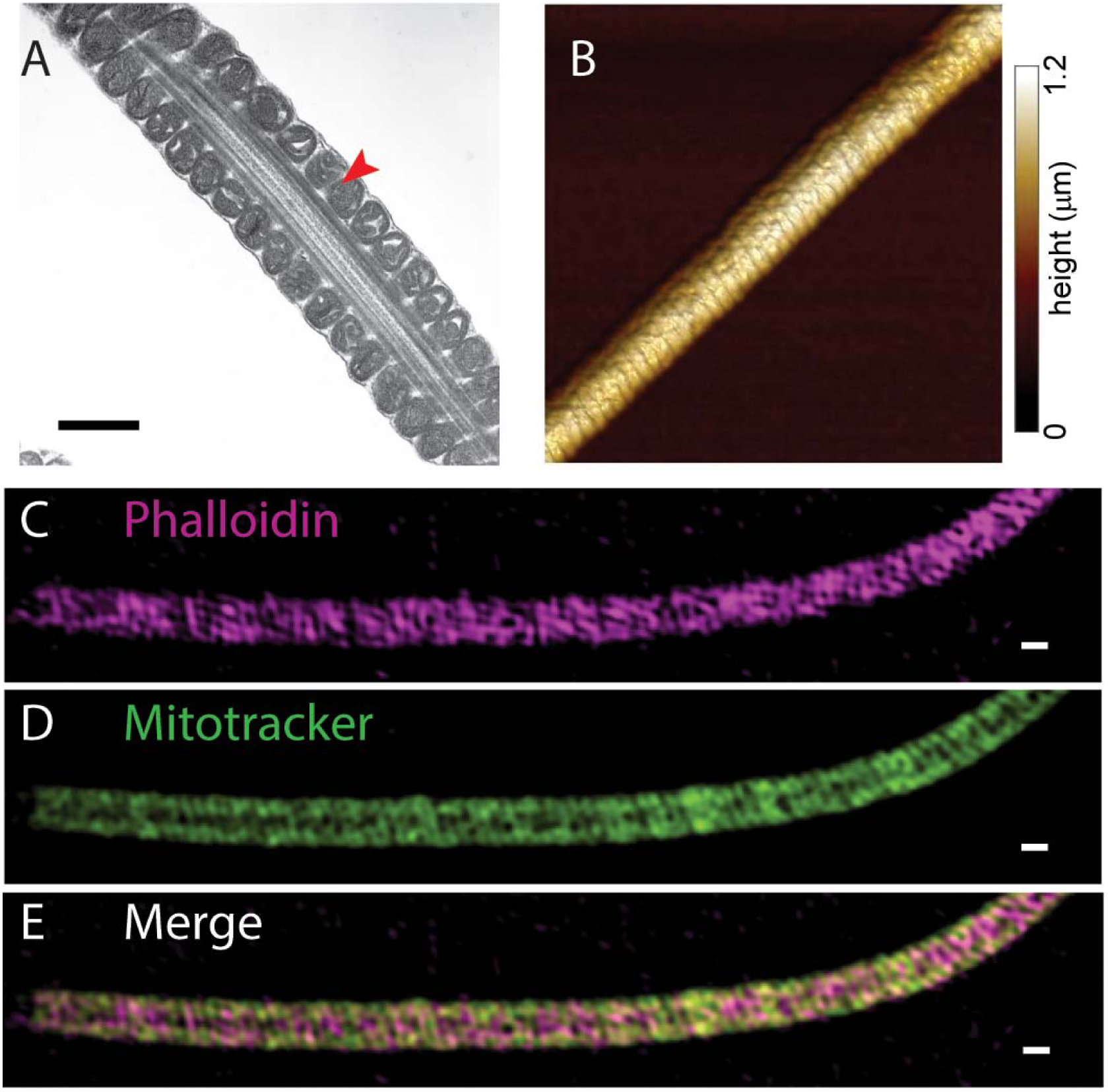
Organization of the sperm midpiece. **(A)** TEM image of the sperm midpiece showing the mitochondria, axoneme, and plasma membrane. Red arrowhead indicates a distance of 300 nm from the center of the flagellum. The scale bar is 500 nm. **(B)** Surface topography of sperm midpiece as obtained by AFM. **(C)** SIM reconstruction of actin labeled with phalloidin-AF647 in the sperm midpiece. **(D)** SIM reconstruction of mitochondria labeled with Mitotracker in the sperm midpiece. **(E)** Merge of phalloidin and mitotracker reconstructions shown in panels C and D. The scale bars in panels C-D are 500 nm.

### Structural actin-associated proteins in the sperm midpiece

Two of the most widespread structural actin-associated proteins are spectrin and adducin. Often, actin, spectrin, and adducin form complex structures (Baines, 2010). Well-known examples are found in erythrocytes (Anong et al., 2009) and neurons (Xu et al., 2013). Immunofluorescence microscopy revealed that both spectrin and adducin, are present in the head and throughout the entire flagellum of mouse sperm (Fig. 4 A and 4 B). A STORM superresolution image of spectrin in a sperm flagellum is shown in Fig. 4C, and a zoom of the midpiece is shown in Fig. 4D. The cross-section of the midpiece showed that spectrin is absent from the central axoneme (Fig. 4 E). The analysis of radial localization indicated a maximum density at 410 ± 23 nm (mean ± sd, n = 4) in the midpiece (Fig. 4 F), which suggests that spectrin is localized apposed to the plasma membrane. Similar results were obtained when sperm were incubated with a different anti-spectrin antibody (Suppl. Fig. 1).

**Figure 4:**
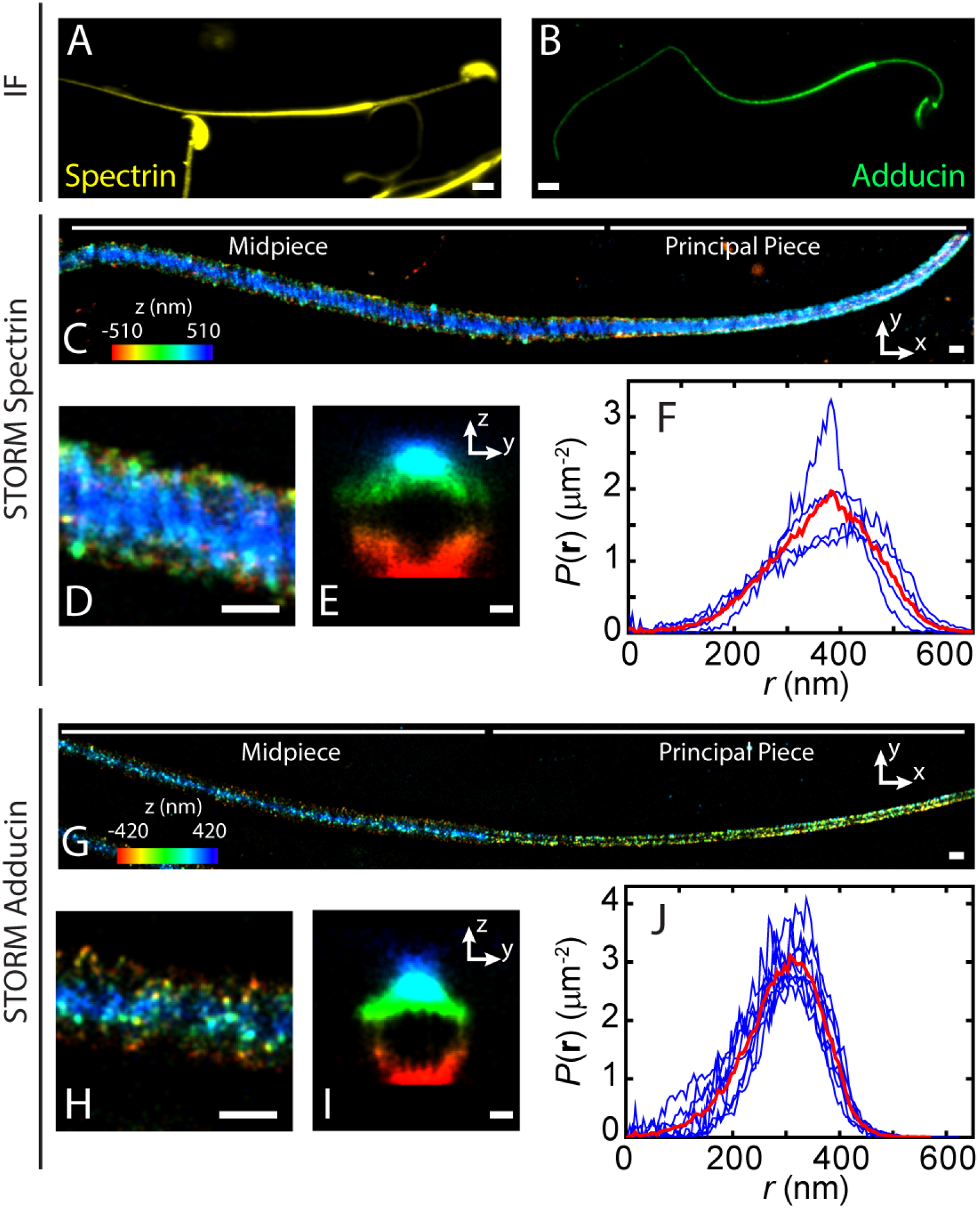
Localization of spectrin and adducin in the sperm. **(A)** Spectrin immunofluorescence (IF) in sperm cells. **(B)** Adducin immunofluorescence (IF) in mouse sperm. **(C)** STORM reconstruction of spectrin in the sperm flagellum where the structure in both the midpiece and principal piece are observed. **(D)** Zoom of a section of the midpiece of panel C. **(E)** Composite sum of intensity projection of spectrin cross-sections along 8.3 μm of the sperm midpiece. **(F)** Radial distribution of spectrin localization in the midpiece. Blue lines are distributions from four individual cells and the thick red line is their average. **(G)** STORM reconstruction of adducin in the sperm flagellum where the structure in both the midpiece and principal piece are observed. **(H)** Zoom of a section of the midpiece of panel G. **(I)** Composite sum of intensity projection of adducin cross-sections along 3.4 μm of the sperm midpiece. **(J)** Radial distribution of adducin localization in the midpiece. Blue lines are distributions from eight individual cells and the thick red line is their average.

STORM images of adducin show that this protein forms a sparse dotted structure in the midpiece and a more uniform structure along the principal piece (Fig. 4 G). A zoom of the midpiece structure is shown in Fig. 4 H. The cross-section of the midpiece indicated that adducin is also absent from the central axoneme (Fig. 4 I). The analysis of radial localization indicated that the highest density of adducin peaks at 306 ± 16 nm (n = 8) in the midpiece. This localization is coincident with the localization of F-actin found in the same region.

### Structure of the actin cytoskeleton and actin-associated proteins in the principal piece

The structural organization of the principal piece is key for sperm function and motility. We evaluated the organization of the actin cytoskeleton and actin-interacting proteins in the principal piece. F-actin formed short bundles distributed throughout the principal piece (Fig. 5 A). The cross-section and the radial distribution analysis indicated that the maximum density of F-actin bundles is next to the axoneme and extended outwards to a radius of 300 nm (Fig. 5 B and C), independent of the distance to the annulus (Fig. 5 D). Spectrin was found uniformly distributed along the length of the principal piece (Fig. 5 E). The cross-section analysis showed that spectrin localizes in a ring-like structure (Fig. 5 F). The radius was found to be largest at the annulus where it is 305 ± 25 nm (Fig. 5 G) and tapered towards the end of the flagellum (Fig. 5 H). This localization was consistent with cortical localization of spectrin, as the sperm flagellum became narrower towards the end piece. Adducin localization was uniform in the principal piece (Fig 5 I). The cross-section showed that this protein also localized in a ring-like structure (Fig 5 J) that extended longitudinally throughout the flagellum. The radial distribution analysis indicated that the maximum density of adducin is at a radius *r* = 140 nm, and that this radius is independent of the distance from the annulus (Figs. 5 K and L).

**Figure 5:**
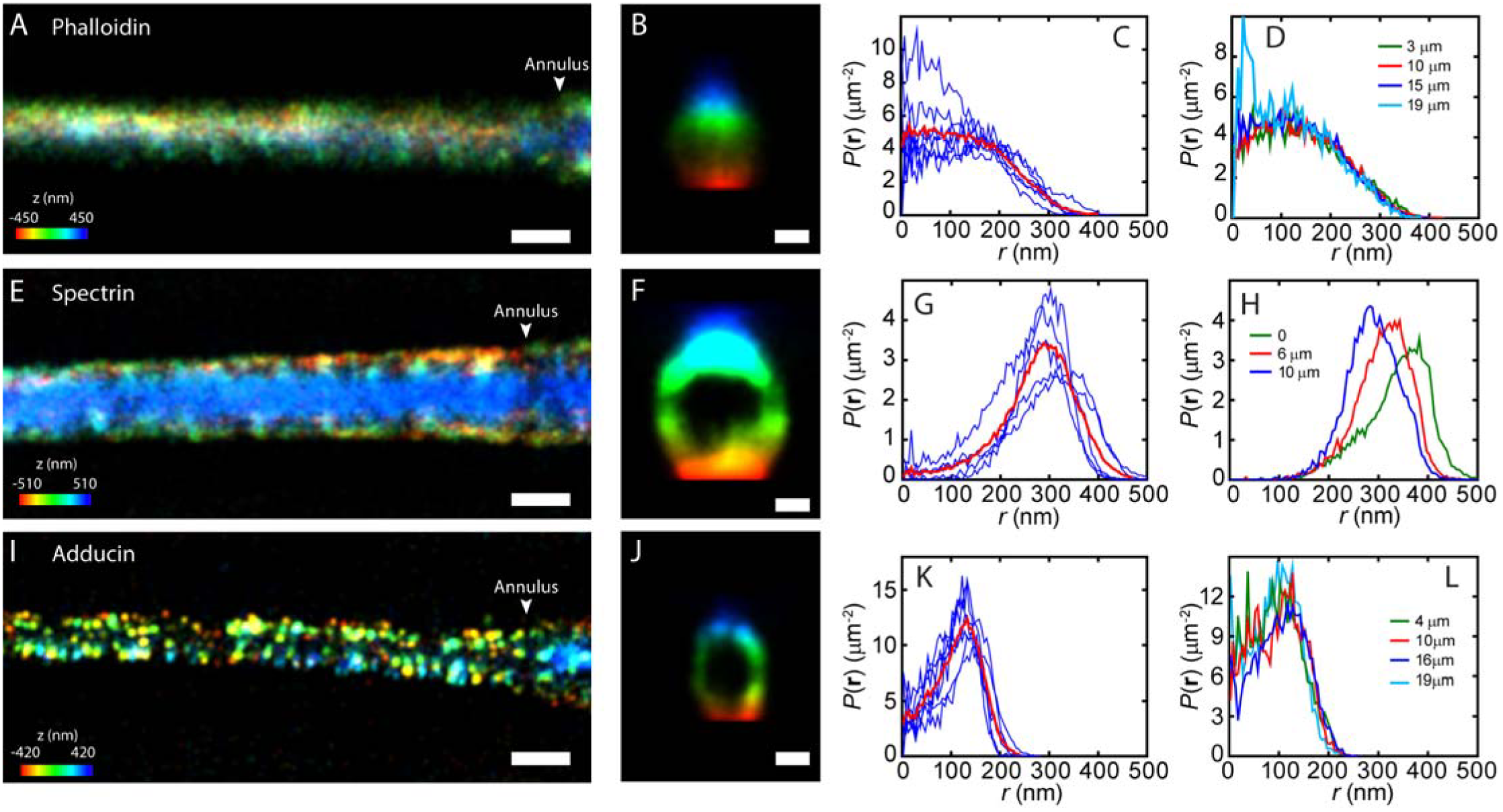
Actin-based cytoskeleton in the principal piece. **(A)** STORM reconstruction of actin labeled with phalloidin-AF647 in the sperm principal piece. Part of the principal piece is observed in the image. The reconstruction is colored according to the localization height as indicated in the colormap. **(B)** Composite sum of intensity projection of actin cross-sections along 3.7 μm of the sperm principal piece. **(C)** Radial distribution of F-actin localization in the principal piece. The blue lines are distributions from seven individual cells and the thick red line is their average. **(D)** Radial distribution of F-actin localization in the principal piece dependent on the distance from the annulus of one representative cell. **(E)** STORM reconstruction of spectrin in the sperm principal piece. **(F)** Composite sum of intensity projection of spectrin cross-sections along 5.1 μm of the sperm principal piece. **(G)** Radial distribution of spectrin localization in the principal piece. Blue lines are distributions from five individual cells and the thick red line is their average. **(H)** Radial distribution of spectrin localization in the principal piece dependent on the distance from the annulus of one representative cell. **(I)** STORM reconstruction of adducin in the sperm principal piece. **(J)** Composite sum of intensity projection of adducin cross-sections along 3.5 μm of the sperm principal piece. **(K)** Radial distribution of adducin localization in the principal piece. Blue lines are distributions from seven individual cells and the thick red line is their average. **(L)** Radial distribution of adducin localization in the principal piece dependent on the distance from the annulus of one representative cell. The scale bars in panels A, E and I are 500 nm, in panels B, F and J are 200 nm.

## Discussion

Similar to other cylindrical biological structures, the sperm flagellum relies on the cytoskeleton for its structural organization and specialized mechanical properties. Along its longitudinal axis, the sperm flagellum is organized in three different sections: the midpiece, the principal piece, and the end piece. We used three-dimension STORM to study the structure of the actin cytoskeleton in the mouse sperm flagellum. Our results indicate that actin filaments and actin-associated proteins are differentially organized in the midpiece and the principal piece.

In the midpiece, mitochondria are organized in a unique helical sheath enclosing the central axoneme and the ODFs (Fawcett, 1975). The proper organization of the mitochondrial sheath is essential for sperm function and fertility (Bouchard et al., 2000; Kissel et al., 2005; Suzuki-Toyota et al., 2004). Defects of sperm mitochondrial ultrastructure have been associated with decreased sperm motility and fertility (Pelliccione et al., 2011). In this work, we show that in the midpiece F-actin forms a double-helical structure with a periodicity of 244 nm. This actin structure is coincident with the organization of the mitochondrial sheath in mouse sperm (Ho and Wey, 2007). Such double-helix structure represents a novel actin architecture that, to the best of our knowledge, has not been previously reported. Interestingly, it has been shown that actin forms helix-like structures in the flagellum of the protozoan intestinal parasite *Giardia intestinalis* (Paredez et al., 2011). These findings open the question whether, to some extent, flagellar helical structures are conserved among diverse species.

Previous works have shown the existence of a dense cytoskeletal structure in the midpiece related to mitochondria, however the composition of this structure has remained unknown (Olson and Winfrey, 1986; Olson and Winfrey, 1990). Our results place actin and its related proteins as potential components of this cytoskeletal structure. In various cell types, actin-dependent immobilization of mitochondria is critical for localization of these organelles at sites of high ATP utilization (Boldogh and Pon, 2006). Our results suggest that F-actin structures in the midpiece are related to the integrity of the mitochondrial sheath and the localization of mitochondria. The actin cytoskeleton could be involved in the migration of mitochondria to the midpiece during spermiogenesis, and in providing a scaffold that confines mitochondria at this cellular compartment. In support of this hypothesis, genetically modified male mice lacking the actininteracting nectin-2 have an unorganized mitochondrial sheath and F-actin is absent from the sperm midpiece (Bouchard et al., 2000). These mice are infertile and, although sperm motility appears normal, long-term periods of incubation in capacitating media showed loss of motility *in vitro* and reduced migration of the sperm into the oviduct *in vivo* (Mueller et al., 2003). These results highlight the importance of the actin cytoskeleton in the midpiece for normal sperm function and fertility.

To have a more comprehensive understanding of the actin cytoskeletal architecture in the midpiece, we evaluated the presence and localization of the actin-associated proteins spectrin and adducin. Spectrin is a protein of high molecular weight that is part of the cortical skeleton (Machnicka et al., 2014). Spectrin reversibly unfolds and refolds when subjected to forces up to 20 pN (Discher and Carl, 2001), acting as a molecular spring and dramatically altering the elasticity of actin-spectrin meshwork. Spectrin has been previously described in sperm of guinea pig and bull where it localizes in the sperm head acrosome and along the flagellum (Hernandez-Gonzalez et al., 2000; Yagi and Paranko, 1995). In the present work, we find that spectrin localizes in the head and flagellum of mouse sperm. In addition, our superresolution imaging indicates that spectrin localizes close to the flagellum plasma membrane. Before acquiring fertilization-competence, sperm change their motility pattern from progressive to hyperactivated motility (Gervasi and Visconti, 2016). Hyperactivation consists in highly asymmetrical waveforms of the tail combined with an increase in the amplitude of the flagellar bending (Olson et al., 2011). Therefore, during hyperactivated motility the sperm midpiece experiences severe bending (Lindemann and Lesich, 2016). Taking into consideration the elastic properties of spectrin, we hypothesize that this protein reinforces the structure of the mitochondrial sheath and gives the midpiece the required elasticity during sperm hyperactivation. Rigid midpieces lead to male infertility due to failure in the acquisition of hyperactivated motility (Miyata et al., 2015), which emphasizes the importance of an elastic midpiece structure for sperm function. Adducin is an actin capping protein that promotes the binding of F-actin to spectrin (Li et al., 1998). We found that adducin is present in mouse sperm and is localized in the acrosome as well as in the tail. Conversely to the F-actin and spectrin structure found in the midpiece, adducin presented a dotted distribution in this region. This discrete localization found at the same radius as F-actin, and close to the radius of spectrin in the midpiece suggests that adducin might be linking the F-actin and spectrin structures observed in the midpiece (see model in Fig. 6 A).

**Figure 6:**
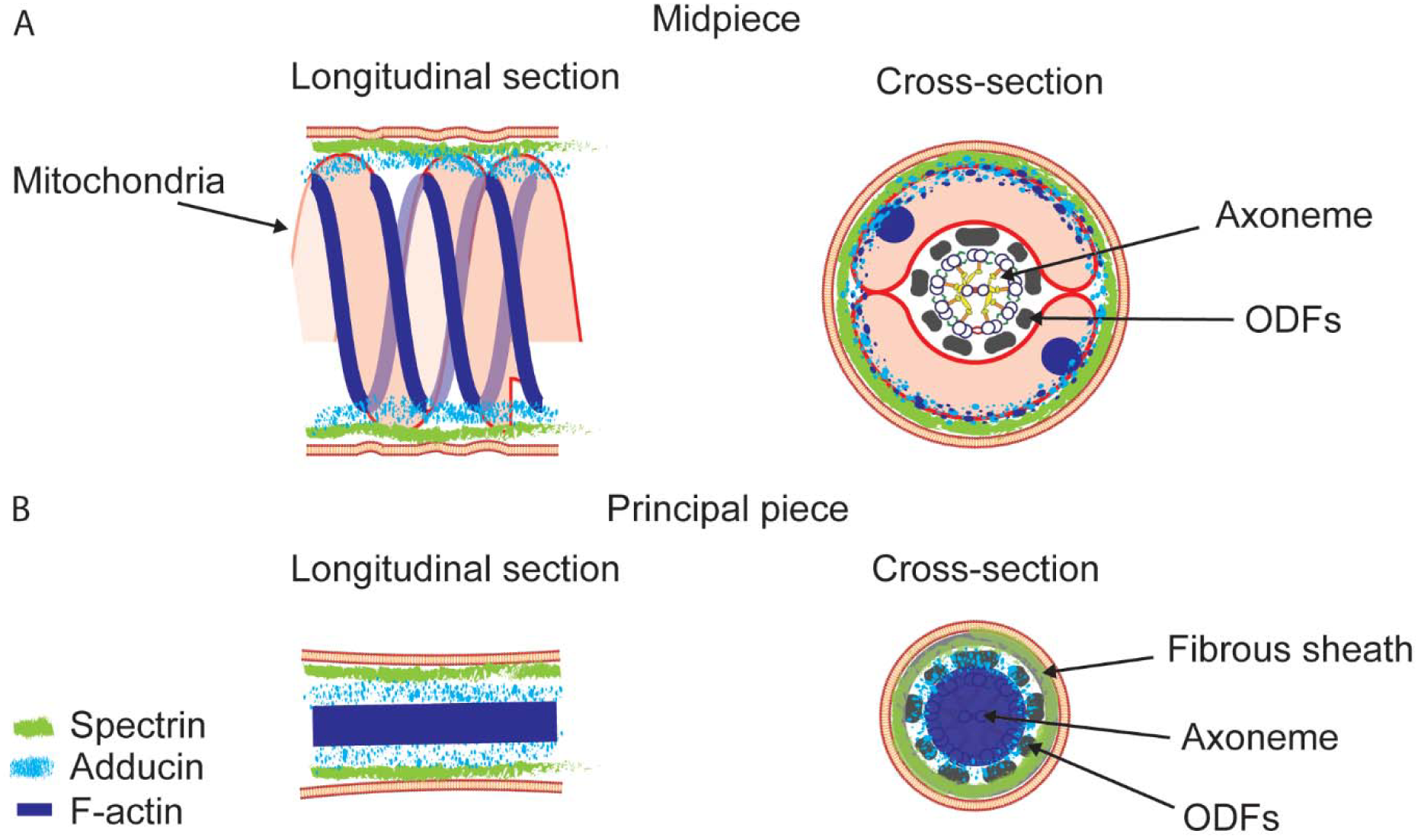
Schematic representation of the actin-based cytoskeleton in the mouse sperm flagellum. **(A)** Sketch of the localization of F-actin (blue), spectrin (green), and adducin (cyan) in a longitudinal section and cross-section of the sperm midpiece. Midpiece structures such as mitochondrial sheath, outer dense fibers (ODFs) and central axoneme are indicated. **(B)** Sketch of the localization of F-actin (blue), spectrin (green), and adducin (cyan) in a longitudinal section and cross-section of the sperm principal piece. Principal piece structures such as fibrous sheath, ODFs and central axoneme are indicated.

In neurons, the cytoskeleton of axons is formed by periodical actin rings that intercalate with spectrin, and sodium channels localize following this underlying actin-spectrin-based cytoskeleton (Xu et al., 2013). The sperm specific Ca^2+^-channel CatSper organizes in four longitudinal backbones along the principal piece (Chung et al., 2017; Chung et al., 2014). Delocalization of these channels results in the impossibility of acquiring hyperactivated motility (Chung et al., 2014). The actin-adducin-spectrin cytoskeleton of the sperm principal piece likely plays a role in maintaining highly organized structures by facilitating the arrangement of specific proteins in organized membrane domains, and thus regulating sperm function (see model in Fig. 6 B). During capacitation, the motility pattern of the sperm changes from progressive to hyperactivated. Structural organization of ion channels and pumps is critical for the modulation of the dynamic processes associated with the acquisition of the asymmetrical hyperactivated motility.

In the same way that erythrocytes remain for long periods of time in the blood stream, sperm are exposed to different environments and can survive for days inside the female reproductive tract prior to fertilization. The structures of F-actin, spectrin, and adducin suggest a role of these proteins in maintaining sperm integrity, elasticity and cell homeostasis. In line with these functions, it has been shown that mouse sperm present resistance to swelling in hypoosmotic media due to F-actin in the sperm tail (Noiles et al., 1997). The actin cytoskeleton of the principal piece could contribute to the resistance to osmotic stress.

Despite the relevance of the actin cytoskeleton in the regulation of cell physiology, its role in sperm function and its relationship with other capacitation-induced signaling processes are not well understood. In the present work, we showed that compartments of the sperm flagellum present different F-actin, adducin and spectrin organization. This highly organized cytoskeleton is expected to play a critical role in sperm function and fertility.

## Methods

### Reagents

Poly-lysine and anti-spectrin antibodies (cat. s-1515, and S-1390) were obtained from Sigma-Aldrich (St Louis, MO). Anti-adducin (cat. ab51130) was purchased from Abcam (Cambridge, UK). Alexa Fluor 488-conjugated phalloidin (cat. A12379), Alexa Fluor 647-conjugated phalloidin (cat. A22287), Alexa Fluor 647-conjugated anti-rabbit secondary antibody (cat.A21244), and MitoTracker Green FM (cat. M7514) were obtained from Thermo Fisher Scientific (Waltham, MA). Paraformaldehyde (cat. 15700) and glutaraldehyde (cat. 16200) were obtained from Electron Microscopy Sciences (Hatfield, PA).

### Animals and sample collection

Young adult (7–8 weeks-old) CD-1 male retired breeders were purchased from Charles River Laboratories (Wilmington, MA). Genetically modified LifeAct-GFP mice were kindly donated by Santiago Di Pietro.

Animals were euthanized in accordance with the Animal Care and Use Committee (IACUC) guidelines of University of Massachusetts Amherst and Colorado State University. Two cauda epididymides were dissected, slightly cut and placed into 1 ml modified Krebs-Ringer medium [TYH’s HEPES-buffered medium with composition: NaCl (100 mM), KCl (4.7 mM), KH_2_PO_4_ (1.2 mM), MgSO_4_ (1.2 mM), glucose (5.5 mM), pyruvic acid (0.8 mM), CaCl_2_ (1.7mM), HEPES (20 mM)]. Sperm were allowed to swim-out for 10 min at 37 °C, and then the epididymides were removed and the suspension of sperm was adjusted to a final concentration of 1–2 10^7^ cells/ml.

### Indirect immunofluorescence

After collection, sperm were centrifuged at 800 g for 5 min, washed with 1 ml PBS, and centrifuged again at 800 g for 5 min. Sperm were fixed in solution by adding 4% (v/v) fresh paraformaldehyde in PBS for 10 min at room temperature. Then sperm were centrifuged at 800 g for 5 min, washed with 1 ml PBS, centrifuged again at 800 g for 5 min and resuspended in 500 μl PBS. Sperm were seeded in polylysinated coverslips (Corning #1.5) and let air-dry. Non-bound cells were removed by washing with PBS. After that, cells were permeabilized with 0.5 % (v/v) Triton X-100 in PBS for 5 min at room temperature and washed 3 times for 5 min with T-PBS (0.1 % (v/v) Tween-20 in PBS). Samples were blocked with 3 % (w/v) BSA in T-PBS for 1 hr at room temperature and then incubated with anti-spectrin antibody s-1515 (1:25), anti-spectrin antibody s-1390 (1:50; supplemental figure 1) or anti-adducin antibody (1:100) diluted in T-PBS containing 1 % (w/v) BSA overnight at 4 °C in a humidifier chamber. After incubation, sperm were washed 3 times for 5 min with T-PBS and incubated with the corresponding Alexa 488-conjugated secondary antibody (1:200) diluted in T-PBS containing 1 % (w/v) BSA for 1 hr at room temperature. Finally, samples were washed 3 times 10 min with T-PBS and the coverslips were mounted using Vectashield H1000 (Molecular Probes, Eugene, OR). Epifluorescence microscopy was performed using a TE300 Eclipse microscope (Nikon) with an oil immersion objective (Nikon, 60x, NA1.4) coupled with a CMOS camera (Zyla 4.2, Andor, Belfast, UK). Negative controls using secondary antibody alone were used to check for specificity.

### STORM sample preparation

For spectrin and adducin labeling, samples were fixed and incubated with primary antibody as described above for indirect immunofluorescence. After primary antibody incubation, cells were washed with T-PBS three times for 5 min each, and then incubated with Alexa Fluor 647-conjugated anti-rabbit secondary antibody diluted in PBS containing 1 % (w/v) BSA (1:1000) for 1 hr at room temperature. Cells were then washed with T-PBS three times for 5 min each. After washing, cells were immediately mounted in STORM imaging buffer (50 mM Tris-HCl (pH 8.0), 10 mM NaCl, 0.56 mg/ml glucose oxidase, 34 μg/ml catalase, 10 % (w/v) glucose, and 1 % (v/v) β-mercaptoethanol).

For phalloidin labeling, sperm samples were seeded in polylysinated coverslips (Corning #1.5) and let air-dry for 15 min. Then, samples were fixed and permeabilized with 0.3 % (v/v) glutaraldehyde - 0.25 % (v/v) Triton X-100 in cytoskeleton buffer [CB, MES (10 mM, pH 6.1), NaCl (150 mM), EGTA (5 mM), glucose (5 mM), and MgCl_2_ (5 mM)] for 1 min. Cells were then washed with CB buffer three times for 5 min each, and then incubated with 2 % (v/v) glutaraldehyde in CB buffer for 15 min. Then cells were washed with CB buffer twice for 10 min each. To avoid background fluorescence caused by glutaraldehyde fixation, samples were incubated with 0.1 % (w/v) of sodium borohydride in PBS for 7 min. After that, samples were washed with PBS twice for 5 min each, and incubated with 0.5 μM of Alexa Fluor 647-conjugated phalloidin in PBS for overnight at 4 °C in a humidifier chamber. Cells were then washed with PBS three times for 5 min each and were mounted in STORM imaging buffer.

### STORM imaging

Images were acquired using Nikon NIS-Elements 4.51 software in a custom-built microscope equipped with an Olympus PlanApo 100x NA1.45 objective and a CRISP ASI autofocus system (Krapf et al., 2016; Weigel et al., 2011). Alexa Fluor 647 was excited with a 638-nm laser (DL638-050, CrystaLaser, Reno, NV) under continuous illumination. Initially the photo-switching rate was sufficient to provide a substantial fluorophore density. However, as fluorophores irreversible photo-bleached, a 405-nm laser was introduced to enhance photo-switching. The intensity of the 405-nm laser was adjusted in the range 0.01 – 0.5 mW to maintain an appropriate density of active fluorophores. Axial localization was achieved via astigmatism using a MicAO 3DSR adaptive optics system (Imagine Optic, Orsay, France) inserted into the emission pathway between the microscope and the EMCCD camera ^(Clouvel et al., 2013; Izeddin et al., 2012)^. This technology achieves three-dimensional localization via shaping the axial point spread function, allowing both the correction of spherical aberrations and the introduction of astigmatism. User-controlled introduction of astigmatism enabled the accurate detection of single-molecules over a thickness of 1 mm and, in turn, 3D reconstruction (Marbouty et al., 2015b) (Marbouty et al., 2015a). A calibration curve for axial localization was generated with 100-nm TetraSpeck microspheres (Invitrogen) immobilized on a coverslip (Huang et al., 2008). The images were acquired in a water-cooled, back-illuminated EMCCD camera (Andor iXon DU-888) operated at -85 °C at a rate of 50 frames/s. 50,000 frames were collected to generate a superresolution image.

### Superresolution image reconstruction

Single-molecule localization, drift correction using image cross correlation and reconstruction were performed with ThunderSTORM (Ovesny et al., 2014).

### Image analysis

To find the molecular radial distributions, we selected regions of interest of the flagellum that were found to lie in a straight line. The center of the flagellar cross section was first found by Gaussian fitting of the localizations histograms along *x* and *y*. The coordinates of the localized molecules were then transformed into cylindrical coordinates to obtain the radial position *r* and azimuth angle *θ*.

### Structured illumination microscopy (SIM)

For dual-color SIM imaging, sperm were labeled in suspension with MitoTracker Green FM (0.2 μM) for 30 min at 37 °C. Sperm were pelleted by centrifugation at 800 g, resuspended in PBS, seeded in polylysinated coverslips (Corning #1.5), and let air dry. Once dried, fixation, permeabilization and staining with Alexa Fluor 647-conjugated phalloidin were performed as described in the STORM sample preparation section above. Images were taken using a Nikon A1R-SIMe microscope using a Nikon PlanApo 100x NA1.49 objective. Image reconstruction was performed in NIS elements software (Nikon).

### Transmission electron microscopy

Luminal fluid collected from cauda epididymides of two mice was pooled and fixed in 2 % (v/v) glutaraldehyde in 0.1 M sodium cacodylate and post-fixed in osmium tetroxide. Samples were embedded in Poly/Bed812 (Polysciences; Warrington, PA). 1-μm-thick sections were cut and stained with toluidine blue for initial light microscopic examination and to locate areas for ultrastructural evaluation. Thin sections (60–80 nm) were cut from the identified areas, stained with uranyl acetate and lead citrate, and evaluated using JEOL JEM 1200EX transmission electron microscope.

### Atomic force microscopy

Sperm cells were placed in polylysinated glass-bottom petri dishes and mounted in PBS. AFM measurements were performed in a Bruker Resolve SPM system using the ScanAsyst mode with a ScanAsyst-Fluid+ probe (Bruker, Billerica, MA).

## Acknowledgements

We thank Santiago Di Pietro for providing the LifeAct mouse. SIM experiments and analysis were performed at the Light Microscopy Facility and Nikon Center of Excellence at the Institute for Applied Life Sciences, UMass Amherst with support from the Massachusetts Life Sciences Center. TEM was performed at ARBL Morphological Services Facility at CSU in consultation with Dr. D. N. Rao Veeramachaneni. We thank Jennifer Palmer and Carol Moeller for TEM technical assistance. We also thank James Bamburg, James K. Graham, and Darío Krapf for useful discussions and Sanaz Sadegh and Patrick Mannion for their help with the experimental setup. This work was supported by the National Science Foundation grant 1401432 (to D.K.), the Eunice Kennedy Shriver National Institute of Child Health and Human Development NIH grants R01HD38082 and R01HD44044 (to P.E.V.), and Fogarty NIH grant R01TW008662 (to M.G.B.).

## Author contributions

M.G.G., P.E.V. and D.K. designed research; M.G.G. and X.X. performed research; M.G.B. and B.C-G. contributed new analytic tools; M.G.G., X.X., and D.K. analyzed data; M.G.G., P.E.V., and D.K. wrote the initial draft.

---

**Supplemental Figure 1:**
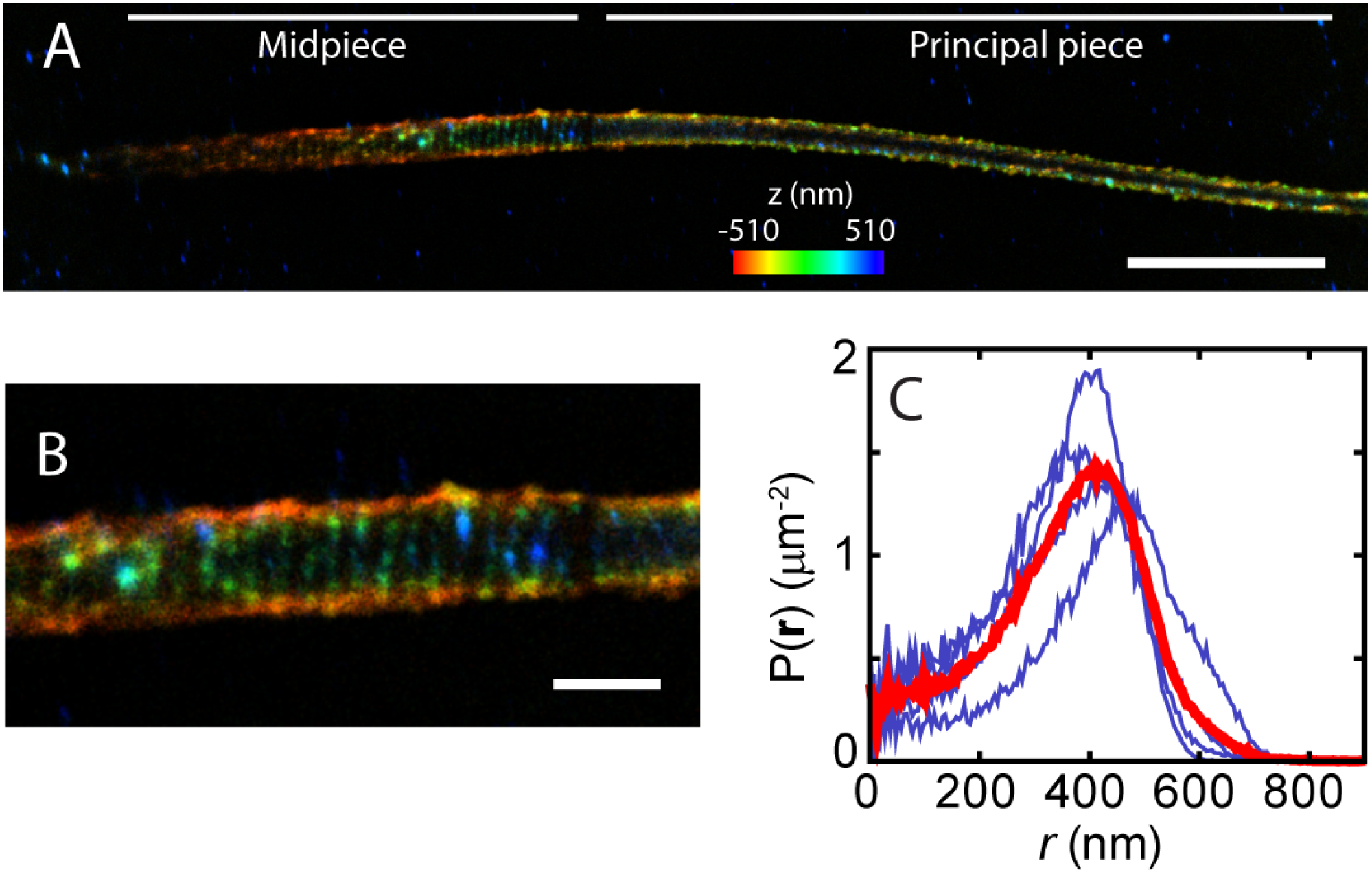
Superresolution localization of spectrin in the sperm midpiece. **(A)** STORM reconstruction of spectrin in the sperm flagellum where the structure in both the midpiece and principal piece are observed. **(B)** Zoom of a section of the midpiece of panel A. **(C)** Radial distribution of spectrin localization in the midpiece. Blue lines are distributions from four individual cells and the thick red line is their average. The scale bar in panel A is 5 μm, and in panel B is 1 μm. Anti-spectrin antibody used was purchased from Sigma (cat. s-1390).

